# Continuous Endoglin (CD105) Overexpression Disrupts Angiogenesis and Facilitates Tumor Cell Metastasis

**DOI:** 10.1101/691824

**Authors:** Claudia Ollauri-Ibáñez, Elena Núñez-Gómez, Cristina Egido-Turrión, Laura Silva-Sousa, Alicia Rodríguez-Barbero, José M. López-Novoa, Miguel Pericacho

**Author notes:** Corresponding Authors Dr. Miguel Pericacho Bustos, University of Salamanca, Edificio Departamental, Campus Miguel de Unamuno, Salamanca, SPAIN 37007, Dr. Elena Núñez Gómez, University of Salamanca, Edificio Departamental, Campus Miguel de Unamuno, Salamanca, SPAIN 37007. These authors have contributed equally to this work (Claudia Ollauri-Ibáñez, Elena Núñez-Gómez).

## Abstract

Angiogenesis is a complex process essential for tumor growth. For this reason, high levels of pro-angiogenic molecules, such as endoglin (CD105), are supposed to be related to greater tumor growth that lead to a poor cancer prognosis. However, we demonstrate here that defects in angiogenesis that can be attributed to high levels of endoglin, lead to development and worsening of cancer disease. Steady endoglin overexpression disrupts the correct stabilization of the endothelium and the recruitment of mural cells. In consequence, endoglin overexpression gives rise to altered vessels that promote the intravasation of tumor cells, the subsequent development of metastases and, thus, a worse cancer prognosis.

## INTRODUCTION

Cancer is a generic term for a large group of diseases characterized by the growth of abnormal cells (tumor cells) beyond their usual boundaries, invading adjoining parts of the body and even spreading to other organs. Tumor tissue is highly proliferative, expansive and has high metabolic requirements; thus, the oxygen demand soon exceeds the input (1). For this reason, as described in the 1970s, solid tumors cannot grow more than a few millimeters in the absence of the development of new blood vessels (2). Angiogenesis is the main event through which these vessels are created. Angiogenesis is the *de novo* formation of blood vessels from preexisting vessels in response to several stimuli, mainly hypoxia. The principal phases of this process include endothelial cell (EC) activation, tip and stalk cell differentiation, basement membrane degradation, EC sprouting and branching, vessel lumen formation, vessel anastomosis and mural cell recruitment, resulting in a new vascular network that provides blood to the hypoxic tissue (3–5). Tumor angiogenesis is characterized by the dysregulation of many of these processes, leading to irregular and chaotic tumor neovessel formation and thus to dysfunctional and more permeable vessels that facilitate the circulation and homing of tumor cells in other tissues (1,6).

Endoglin (CD105) is an auxiliary receptor for several members of the TFG-β superfamily of cytokines. There is a substantial amount of evidence that relates endoglin to angiogenesis. On the one hand, endoglin expression is increased in the endothelium during active angiogenesis (7), specifically at the angiogenic edge, where sprouting takes place (8,9). On the other hand, deficiency in endoglin expression is responsible for hereditary hemorrhagic telangiectasia type-1 (HHT-1), a disease characterized by vascular malformations (10). In experimental models, the lack of or deficiency in endoglin produces minor and defective angiogenesis (8,11–13).

Because angiogenesis is essential for the growth of tumors, the quantification of microvessel density is a frequent clinical practice to assess tumor prognosis (14–16). More specifically, a high number of endoglin-positive microvessels detected by immunohistochemistry has been associated with poor prognosis in some solid tumors (17,18). Moreover, enhanced endoglin expression in tumor microvessels is a better outcome predictor for some cancer types than the levels of other angiogenesis-related molecules (19,20). In addition, treatment with the anti-endoglin monoclonal antibody TRC105 or with different anti-endoglin shRNAs, siRNAs or miRNAs reduces tumor growth through the reduction of blood vessels inside the tumor (21). All these observations have led to the assumption that the worse prognosis of tumors with high levels of endoglin is due to greater angiogenesis and, therefore, greater tumor growth, although this hypothesis has not been experimentally demonstrated. Recently, it has been shown that anti-endoglin therapy reduces the generation of metastases (22,23); however, the exact mechanisms by which endoglin could contribute to the generation of metastases have not been reported.

Here, we demonstrate that continuous endoglin overexpression does not enhance tumor growth or vascularization. However, it produces alterations throughout the course of angiogenesis that prevent blood vessels from maturing and stabilizing, leading to even poorer quality tumor vessels and facilitating the intravasation and metastasis of tumor cells. For this purpose, we used transgenic mice overexpressing endoglin (*ENG*^*+*^) in a C57BL/6J background, with C57BL/6J mice as controls (WT) (24), and several *in vitro* approaches to study angiogenesis and the specific events that take place during blood vessel formation.

## MATERIALS AND METHODS

### Mice

All animal procedures were European Community Council Directive (2010/63/EU) and Spanish legislation (RD1201/2005 and RD53/2013) compliant and approved by the University of Salamanca Ethical Committee. Animal selection was genotype based and no randomization or blinding was performed. Animals were housed under specific pathogen-free conditions in University of Salamanca facilities (ES-119-002001 SEARMG), in a temperature-controlled room with 12-hour light/dark cycle and reared on standard chow and water provided *ad libitum*. The generation and characteristics of the transgenic mice ubiquitously overexpressing endoglin (*ENG*^*+*^) are described in a previous article from our group (24). Wild-type C57BL/6J mice (WT) were used as controls. For animal anesthesia, 2% isoflurane in oxygen was used. During recovery from the anesthesia, heat was provided and, when necessary, a dose of buprenorphine (0.05 mg/kg) was subcutaneously administered. Animals were sacrificed by CO_2_ inhalation or cervical dislocation, depending on the age and the experiment requirements.

### Cells and cell cultures

LLC cells, provided by Dr. Aliño (Department of Pharmacology, University of Valencia, Spain), were cultured in Dulbecco’s modified Eagle’s medium (DMEM) (Thermo Fisher Scientific) supplemented with 10% fetal bovine serum (FBS) (Thermo Fisher Scientific) and 50 U/mL penicillin-streptomycin (Thermo Fisher Scientific). GFP-LLC cells were generated following “Cell infection” (<u>dx.doi.org/10.17504/protocols.io.23mggk6</u>). MLECs were isolated and purified following “Mouse Lung Endothelial Cell (MLEC) culture” (<u>dx.doi.org/10.17504/protocols.io.28ighue</u>). The human EC line EA.hy926 was supplied by the ATCC and cultured in DMEM, 10% FBS and 50 U/mL penicillin-streptomycin. EA.hy926 cells were stably infected with a vector containing the sequence encoding human endoglin (*ENG*^*+*^) or an empty vector (*Mock*), according to the above-mentioned cell infection protocol. HBVPs were supplied by ScienCell™ and cultured in pericyte medium (PM), 2% FBS, 1% pericyte growth supplement and 1% penicillin/streptomycin (ScienCell™). HBVPs were cultured in petri dishes coated with 2 μg/cm^2^ of poly-L-lysine (Sigma-Aldrich). All cell lines were maintained in 90% RH, 5% CO_2_ atmosphere at 37 °C.

### Tumor angiogenesis assays

Tumors were generated following a LLC xenograft model (<u>dx.doi.org/10.17504/protocols.io.ykkfuuw</u>). Ten days after the injection tumor hemoglobin and DNA content was evaluated (<u>dx.doi.org/10.17504/protocols.io.28jghun</u>). Lung metastases and circulating tumor cell (<u>dx.doi.org/10.17504/protocols.io.28kghuw</u>) were quantified in mice injected with GFP-LLCs.

### *In vivo* angiogenesis assays

Mouse hindlimb ischemia was performed (<u>dx.doi.org/10.17504/protocols.io.28nghve</u>) and perfusion was evaluated in the ischemic and non-ischemic limbs using Doppler Laser Moor LDLS (Moor Instruments) on days 1, 3, 5, 7, 14, 21 and 28 after ischemia. For histological analysis, the soleus muscles were collected 14 days after surgery. Direct *In Vivo* Angiogenesis Assay (DIVAA™) (<u>dx.doi.org/10.17504/protocols.io.28pghvn</u>) was used for quantification of the blood-invaded distance and the content in endothelial cells, while its analog “Plugs of Matrigel^®^” (<u>dx.doi.org/10.17504/protocols.io.28qghvw</u>) was used for RNA isolation and histological studies. Retinas from P6 and P17 were dissected (<u>dx.doi.org/10.17504/protocols.io.28rghv6</u>) and vascularization was evaluated by immunofluorescence and by mural and endothelial marker expression. Immunofluorescence antibodies: FITC-lectin B4 (Ref. 3450-048-FL; Trevigen), anti-VE-cadherin (Ref. 555289; BD Pharmingen), anti-Ki67 (Ref. Ab15580; Abcam), and anti-NG2 (Ref. AB5320; Merck). Three-dimensional rendering of confocal Z-stacks was performed with Fiji software. The areas occupied by the pericytes were quantified using Adobe Photoshop CS6. The “patching algorithm” for MATLAB, provided by Dr. Bentley (Department of Immunology, Genetics and Pathology, Uppsala University, Sweden), was used for VE-cadherin patterning quantification. Sprouting was also evaluated by aortic ring assay (<u>dx.doi.org/10.17504/protocols.io.28sghwe</u>).

### *In vitro* angiogenesis assays

Endothelial cell organization was evaluated by capillary-like structures in Matrigel^®^ assay (<u>dx.doi.org/10.17504/protocols.io.28ughww</u>). Moreover, pericyte incorporation to capillary-like structures (<u>dx.doi.org/10.17504/protocols.io.28vghw6</u>) and pericyte adhesion to an endothelial monolayer (<u>dx.doi.org/10.17504/protocols.io.28wghxe</u>) were assessed. Cell stability was also evaluated *in vitro.* For this aim, two 35-mm culture dishes were seeded with 3×10^4^*ENG*^*+*^ or WT MLECs. Cells from one dish were collected for RNA isolation after 8 hours; cells from the other dish were cultured up to 100% confluence prior to RNA isolation.

### Endothelial cell proliferation and migration

Endothelial cell proliferation was evaluated by “direct cell proliferation assay” (<u>dx.doi.org/10.17504/protocols.io.28xghxn</u>). Moreover, the BrdU incorporation ELISA kit (Ref. 11647229; Roche) was used following manufacturer’s instructions. 10^3^ MLEC or EA.hy926 ECs per well were plated in 96-well plates. After 24 hours, cells were incubated with BrdU (4 hours for EA.hy926 ECs and 24 hours for MLECs). 100 μL of substrate solution was added and, after 5 to 30 minutes, fluorescence at 370 nm was read. Cell motility was assessed following “Wound healing migration assay (Scratch assay)” (<u>dx.doi.org/10.17504/protocols.io.28yghxw</u>), while migration and ECM invasion were assessed following “Transwell assay” (<u>dx.doi.org/10.17504/protocols.io.28zghx6</u>).

### Histological tissue analysis

Tumors and limbs (soleus muscle) intended for histological analysis were fixed in 4% PFA for 24 hours, dissected and embedded in paraffin. 3-μm sections were stained by hematoxylin-eosin (standard protocol) or processed for immunohistochemistry. Immunohistochemistry was carried out using a DISCOVERY ULTRA system (Roche) and the DISCOVERY ChromoMab DAB kit (RUO) (Ref. 760-159, Roche). Sections were incubated with the primary antibody anti-Pecam1 (Ref. ab28364, Abcam) for 1 hour and with OmniMap™ Anti-Rb HRP (Ref. 760-4311, Roche) for 12 minutes. For tumors, 20 images per slide were analyzed for the area occupied by erythrocytes and the number of vessels using Fiji software. For muscles, 25 to 30 images per slide were analyzed for the number of vessels, vessel size and Pecam-1-positive area with Fiji.

### Protein expression

Endoglin expression was evaluated in mouse tissue and in EA.hy926 cells and MLECs by Western blot (<u>dx.doi.org/10.17504/protocols.io.282ghye</u>). Antibodies: anti-human endoglin (Ref. H300; Thermo Fisher Scientific) and anti-mouse β-actin (Ref. A5441; Sigma-Aldrich). Endoglin expression was also evaluated in endothelial cells by flow cytometry (<u>dx.doi.org/10.17504/protocols.io.283ghyn</u>) and by immunofluorescence (<u>dx.doi.org/10.17504/protocols.io.284ghyw</u>). For flow cytometry, the monoclonal antibody PE mouse anti-human CD105 (Ref. 560839, BD Biosciences) was used. Cells were acquired though FACSCalibur™ (BD Biosciences) and results were analyzed using Infinicyt™ software (Cytognos). For immunofluorescence, the anti-human endoglin hybridoma TEA1/58.1 (provided by Dr. Sánchez-Madrid, CNIC, Spain) and secondary antibody anti-muse IgG AlexaFluor568^®^ (Ref. A10037, Thermo Fisher Scientific) were used.

### Gene expression

RNA from tumors, cultured cells, plugs of Matrigel^®^ and retinas were isolated using NucleoSpin^®^ RNA (Ref. 740955; Macherey-Nagel) according to manufacturer’s instructions. RNA from sprouts of aortic rings was extracted using NucleoSpin^®^ RNA XS (Ref. 740.902; Macherey-Nagel). In samples with high Matrigel^®^ content (plugs and sprouts) manufacturer’s instructions for fibrous tissues were followed. cDNA synthesis was performed with iScript RT Supermix (Ref. 170-8841; Bio-Rad). cDNA from sprouts and plugs was preamplified with ssoAdvanced PreAmp Supermix (Ref. 1725160; Bio-Rad) and Prime PCR Preamp Assays 2X (Bio-Rad), listed in Supplementary Table 1. For qPCR, Supermix iQTM SYBR^®^ Green (Ref. 170-8882; Bio-Rad) and an iQTM 5 System (Bio-Rad) were used. Gene expression results were normalized to ribosomal protein S13 (Rps13) and β-actin (Actb) for mouse genes, to glycealdehyde-3-phosphate dehydrogenase (Gapdh) for human genes and to EC markers when we approximated the gene expression in EC content within the tissue. The primers used are listed in Supplementary Tables 1 and 2.

### Statistical analysis

Results (except qPCR) are expressed as the mean ± SEM. qPCR results are represented in box plots that show the median and the 25^th^-75^th^ percentiles, with whiskers showing the 10^th^-90^th^ percentiles. For the *in vivo* experiments, at least 8 animals were used per group, when possible. *In vitro* experiments were repeated at least three times. The D’Agostino-Pearson normality test was applied prior to statistical comparison (Kolmogorov-Smirnov test was used for small datasets). For normally distributed datasets, Student’s T-test was used (Mann-Whitney U-test was used as nonparametric test). Two-way repeated measures ANOVA was used to assess in time-course experiments. Sidak’s post-hoc test was used after ANOVA. Analyses were performed with GraphPad 6 software.

## RESULTS

### Constitutive endoglin overexpression increases the tortuousness of tumor vessels

Subcutaneous injection of Lewis lung carcinoma (LLC) cells in *ENG*^*+*^ and WT mice results in the generation of solid tumors in the flanks of the mice. Contrary to what we expected, the tumors that developed in *ENG*^*+*^ mice did not grow larger than those in the WT mice; no significant differences were found between the weight of the tumors in the two mouse lines (Fig. 1a). Moreover, immunohistochemistry of Pecam1 did not show differences in the number of vessels generated inside the tumors (Fig. 1b-c). Accordingly, the expression levels of murine endothelial markers such as endogenous endoglin (*Eng*) and *Pecam1* were analyzed in tumors by qPCR, and no differences were found between *ENG*^*+*^ and WT tumors (Supplementary Fig. 1a). However, hematoxylin/eosin staining showed that the vessels inside the tumors developed in *ENG*^*+*^ mice were even more impaired and tortuous than those that developed in WT mice, resulting in increased blood extravasation, blood islets and tumor edema (Fig. 1c-d). This result was confirmed by measuring the hemoglobin concentration in the tumor tissue, which was higher in the tumors from *ENG*^*+*^ than in those from WT mice (Fig. 1e). In summary, although dilated and tortuous vessels are characteristic of tumor vasculature, constitutive overexpression of endoglin enhances this characteristic, presumably by affecting the quality of angiogenesis.

**Fig. 1.**
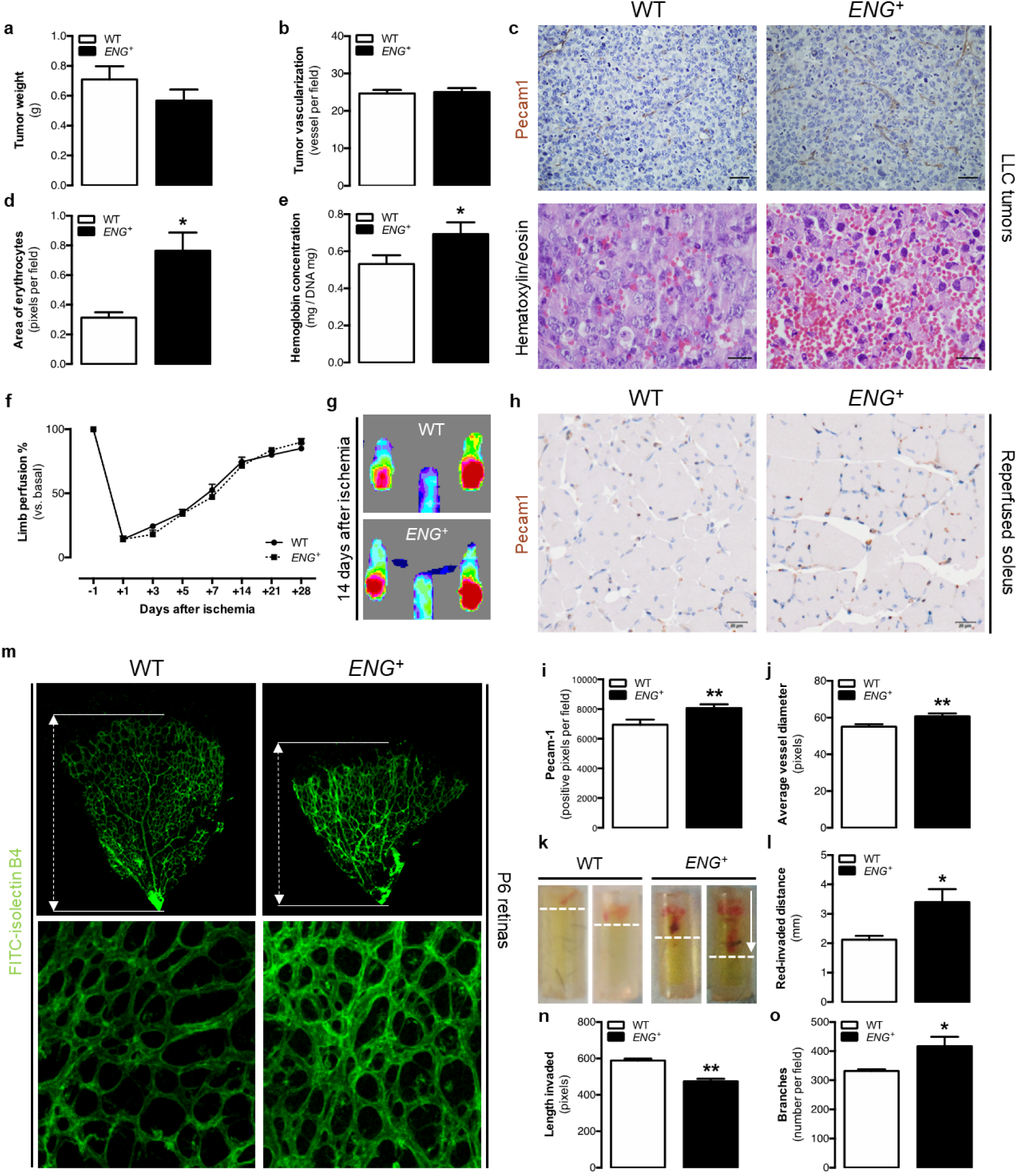
Continuous endoglin overexpression impairs tumor and physiological angiogenesis. (a) Weight of tumors implanted in mice after 10 days [n(WT)=35, n(*ENG*^*+*^)=26; p=0.3329]. (b) Quantification of the number of Pecam1-positive vessels in tumor tissue [n(WT)=4, n(*ENG*^*+*^)=6; p=0.8758]. (c) Upper panel: Pecam1 immunostaining in the tumor tissue showing tumor vessels. Lower panel: Hematoxylin-eosin staining of tumor tissue showing characteristic blood extravasation, blood isles and edema. (d) Quantification of the area covered by erythrocytes in the tumor revealed by hematoxylin-eosin staining [n(WT)=4, n(*ENG*^*+*^)=6; p=0.0484]. (e) Quantification of hemoglobin concentration in the tumor tissue [n(WT)=30, n(*ENG*^*+*^)=21; p=0.0427]. (f) Ratio of ischemic to nonischemic limb perfusion following femoral artery ligation as measured by laser Doppler flow analysis in mice, represented as the percentage of the basal value (before artery ligation) at 1, 3, 5, 7, 14, 21 and 28 days post-ischemia [n(WT)=9, n(*ENG*^*+*^)=9; p=0.4989]. (g) Laser Doppler images showing mice hindlimb perfusion 14 days after ischemia. (h) Pecam1 immunostaining in the ischemic soleus muscle 14 days post-ischemia. (i) Quantification of the Pecam1 immunostained area in the ischemic soleus muscle [n(WT)=3, n(*ENG*^*+*^)=3; p-value=0.0079]. (j) Average Pecam1-positive vessel diameter in the ischemic soleus muscle [n(WT)=3, n(*ENG*^*+*^)=3; p=0.0098]. (k) Angioreactors 9 days after implantation in mice, showing blood invasion. (l) Quantification of angioreactors red-invaded distance from the tube end 9 days after implantation [n(WT)=16, n(*ENG*^*+*^)=28; p=0.0383]. (m) Upper panel: FITC-lectin staining of ECs in the retinal vasculature of p6 pups and representation of the plexus progression. Lower panel: Structure of the vessel plexus of the retina in P6 pups. (n) Quantification of plexus progression in the retinas of P6 pups [n(WT)=7, n(*ENG*^*+*^)=3; p<0.0001]. (o) Quantification of the ramification of the retinas of P6 pups [n(WT)=7, n(*ENG*^*+*^)=3; p=0.0227].

### Sustained endoglin overexpression does not improve post-ischemic reperfusion

To verify whether endoglin overexpression affects the angiogenesis rate, its effect on post-ischemic reperfusion was analyzed. Blood flow in the lower limbs of mice after femoral arterial ligation was measured periodically for 28 days. No significant differences in the reperfusion rates were detected between *ENG*^*+*^ and WT mice throughout this period (Fig. 1f-g). Furthermore, the structure of the blood vessels was assessed in the soleus muscle after 14 days of ischemia by immunohistochemistry of Pecam1. No differences in vessel number were observed between *ENG*^*+*^ and WT mice, but an increased Pecam1-positive area and dilated vessels observed in *ENG*^*+*^ mice indicated the presence of alterations in the vessel structure in the reperfused muscle tissue (Fig. 1h-j).

### Constitutive overexpression of endoglin leads to impaired physiological angiogenesis and an altered structure of the new blood vessels

The direct *in vivo* angiogenesis assay (DIVAA™) consists of the subcutaneous implantation of silicon tubes filled with Matrigel^®^ plus proangiogenic factors into mice. These tubes are invaded by vessels from the mice due to an angiogenic response. The measured distance reached by the red front in each tube was higher in the tubes implanted in *ENG*^*+*^ mice than in those implanted in control mice (Fig. 1k-l), which may suggest increased angiogenesis. However, constitutive endoglin overexpression led to a reduced EC content within the tubes 9 days after implantation (Supplementary Fig. 1d). qPCR analysis of the expression levels of a specific EC marker in the plugs of Matrigel^®^ (analog to DIVAA™) showed no significant differences between *ENG*^*+*^ and WT mice (Supplementary Fig. 1c). The fact that in *ENG*^*+*^ mice, the vessels reach a longer distance with no increase in the number of ECs suggests that the red front observed in *ENG*^*+*^ tubes was not the length invaded by blood-filled vessels but rather extravasated erythrocytes from the defective vessels that moved into the Matrigel^®^, as shown for the tumors.

Another frequently used method to study physiological angiogenesis is the analysis of the development of the retinal vasculature, which takes place between postnatal day 0 (P0) and P8 (25). Retinas were isolated from P6 *ENG*^*+*^ and WT pups, and the vessels were identified with FITC-isolectin B4. Structural analysis revealed that compared with WT retinas, *ENG*^*+*^ retinas showed a lower degree of vascular plexus progression (Fig. 1m-n) and a higher vessel density due to increased ramification (Fig. 1m-o). No differences in the expression levels of the EC markers *Eng* and *Pecam1* were found between retinas from *ENG*^*+*^ and WT mice (Supplementary Fig. 1b), suggesting that the lower degree of plexus progression is not due to a reduced number of ECs but rather to alterations in angiogenesis-driven vascularization.

The effect of endoglin overexpression was also evaluated using the *in vitro* angiogenesis assay. The human EC line EA.hy926 infected with a vector containing human endoglin gene (*ENG*^*+*^) or with the empty vector (Mock) (Supplementary Fig. 2a-b, d-f) was used for this assay, in which the cells spontaneously organize to form capillary-like structures in Matrigel^®^. We found no differences in the number of branches or in the average branch length in the structures created by Mock and *ENG*^*+*^ cells (Supplementary Fig. 2g-i). Thus, it can be concluded that the alterations in angiogenesis observed in tumors and the physiological vascularization demonstrated in *ENG*^*+*^ mice are not due to the inability of ECs to organize themselves into three-dimensional structures.

### An excess of endoglin in ECs reduces proliferation while promoting migration and extracellular matrix invasion

After demonstrating that continuous endoglin overexpression does not affect the ability of ECs to create three-dimensional structures, we studied whether high levels of endoglin alter any of the cellular processes involved in angiogenesis. EC proliferation and migration towards angiogenic stimuli are key highly regulated events during sprouting angiogenesis (26). The effects of endoglin overexpression on these cellular events were assessed *in vitro* using EA.hy926 ECs (*ENG*^*+*^ and Mock) and mouse lung endothelial cells (MLECs) isolated from *ENG*^*+*^ and WT mice (Supplementary Fig. 2c-f).

We studied bromodeoxyuridine (BrdU) incorporation in MLECs and EA.hy926 ECs as an assessment of cell proliferation. The level of BrdU incorporation was significantly lower in *ENG*^*+*^ MLECs and *ENG*^*+*^ EA.hy926 ECs than in WT and Mock cells, respectively (Fig. 2a-b). A tendency to reduce proliferation in *ENG*^*+*^ EA.hy926 ECs compared to Mock cells was also observed by direct cell counts (Fig. 2c).

**Fig. 2.**
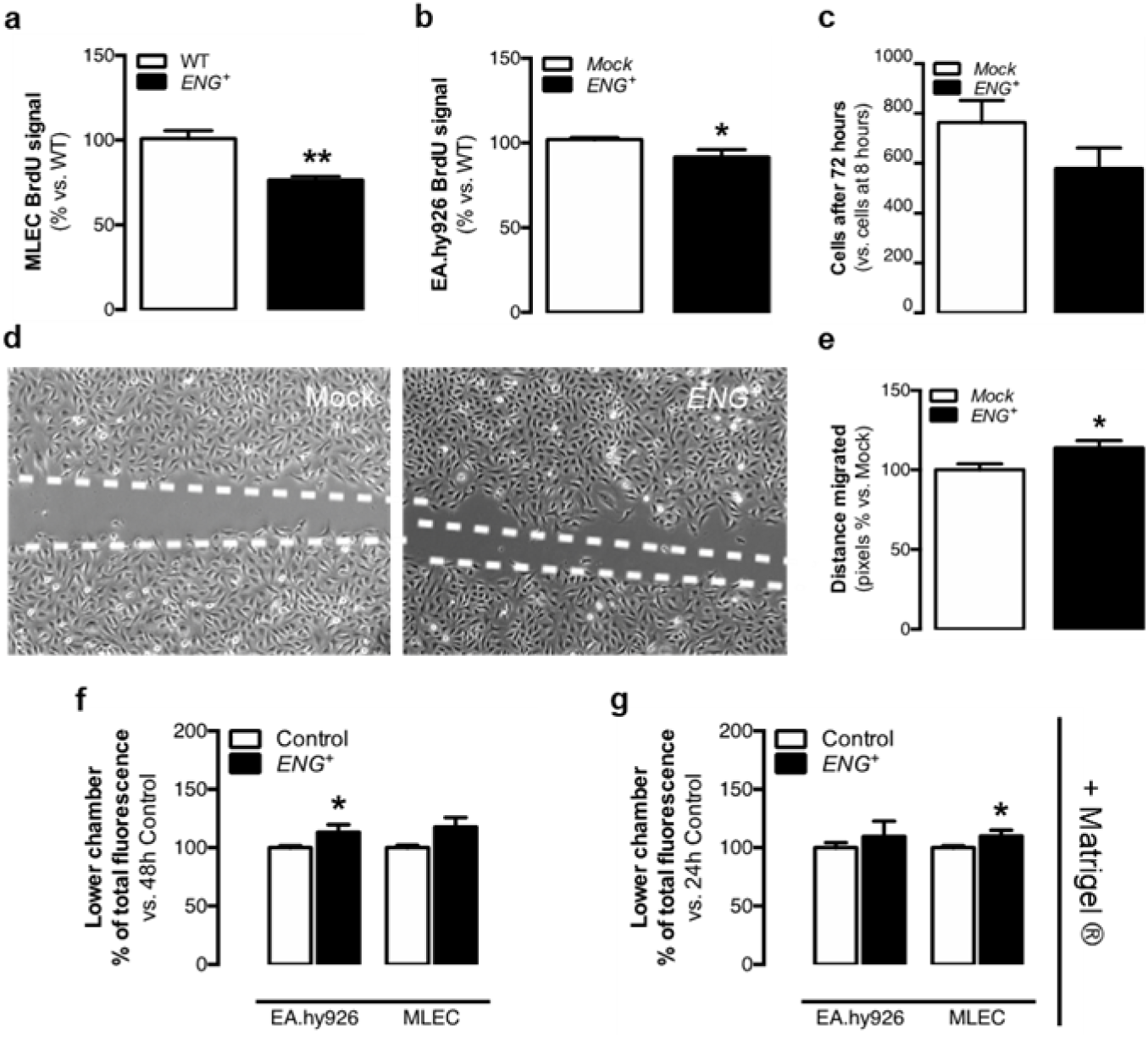
Permanent endoglin overexpression impairs EC physiology *in vitro*. (a) BrdU incorporation after 24 hours in MLECs [n(WT)=3, n(*ENG*^*+*^)=3; p=0.0277]. (b) BrdU incorporation after 4 hours in EA.hy926 cells [n(Mock)=3, n(*ENG*^*+*^)=3; p=0.0272]. (c) Ratio of EA.hy926 cell counts after 72 hours vs. after 8 hours in culture [n(Mock)=3, n(*ENG*^*+*^)=3; p=0.2030]. (d) EA.hy926 scratch closure after 14.5 hours in culture. (e) Quantification of the distance migrated by EA.hy926 cells though the scratch after 14.5 hours [n(Mock)=3, n(*ENG*^*+*^)=3; p=0.0250]. (f) Quantification of EC migration through the uncoated transwell with a VEGF gradient after 48 hours for EA.hy926 and MLEC cells [n(Mock)=3, n(*ENG*^*+*^)=3; n(WT)=3, n(*ENG*^*+*^)=3; p (EA.hy926)=0.0297, p (MLEC)=0.0642]. (g) Quantification of EC migration through the Matrigel^®^-coated transwell with a VEGF gradient after 48 hours for EA.hy926 and MLEC cells [n(Mock)=3, n(*ENG*^*+*^)=3; n(WT)=3, n(*ENG*^*+*^)=3; p (EA.hy926)=0.4453, p (MLEC)=0.0344]. All images shown are representative, and the data are the mean ± SEM.

Cell motility was analyzed by the scratch assay in EA.hy926 EC monolayers in culture. The results showed that the overexpression of endoglin in human ECs increased their motility in culture (Fig. 2d-e). Moreover, we analyzed the ability of EA.hy926 EC and MLEC to migrate towards the angiogenic stimulus VEGF using uncoated transwells and EC invasiveness through the extracellular matrix (ECM) using Matrigel^®^-coated transwells. Enhanced endoglin expression is associated with an increase in cell migration towards VEGF in both EA.hy926 ECs and MLECs (Fig. 2f). Similarly, endoglin overexpression increases ECM invasion in ECs (Fig. 2g).

### Alterations in proliferation and migration caused by endoglin overexpression are not due to modifications in the tip/stalk selection during sprouting

The phenotype of a nonproliferative and highly migratory cell is typical of the so-called tip cells, which are responsible for sprout management during angiogenesis (26). The fact that overexpression of endoglin is associated with less proliferation and more migration and invasion could suggest that endoglin overexpression increases the number of tip cells. To study the first steps of sprouting, we performed an *ex vivo* aortic ring angiogenesis assay with samples from *ENG*^*+*^ and WT mice; in this assay, sprouting from the ring endothelium is induced when it is cultured under proangiogenic conditions. No differences were found in the volume occupied by the sprouts around *ENG*^*+*^ and WT rings on day 5 of the assay (Supplementary Fig. 3a-b). Moreover, the expression levels of endothelial (*Pecam1* and endogen *Eng*) and sprouting markers (*Kdr, Notch1, Dll4* and *Jag1*) (26,27) in the sprout cells were not different between rings from the two mice lines (Supplementary Fig. 3c-d).

We also analyzed the expression of sprouting markers in retina and in plug assay models. Relativization of *Kdr* and *Notch1* expression to the expression of EC markers (*Eng/Pecam1*) in these tissue samples corrects potential differences in EC content. This analysis revealed no differences in *ENG*^*+*^ retinas when compared to WT retinas (Supplementary Fig. 3e). Furthermore, no differences were found between the two mouse lines regarding the expression levels of *Kdr, Notch1, Dll4* and *Jag1* in the plugs (Supplementary Fig. 3f).

These results suggest that, whereas ECs overexpressing endoglin may have a tip-like phenotype, increased levels of endoglin do not affect Notch-induced tip/stalk cell selection or the EC genetic profile.

### Alterations in angiogenesis caused by continuous endoglin overexpression are due to overactive endothelium

We aimed to assess whether altered vascular architecture in mice with constitutive endoglin overexpression is due to an effect on endothelium stability. We analyzed the expression of VE-cadherin (*Cdh5*) in confluent and nonconfluent MLECs. These cells create stable unions when they are confluent, and *Cdh5* gene expression levels directly affect these junctions (28,29). We observed an increase in *Cdh5* expression in confluent WT MLECs compared to the level in nonconfluent cells. This increase was not observed in *ENG*^*+*^ MLECs (Fig. 3a), suggesting a reduction in endothelium stabilization *in vitro*. We also analyzed the gene expression levels of *Cdh5* and the angiopoietin receptor Tie2 (*Tek*), the levels of which are increased in the quiescent endothelium (30), retinas and Matrigel^®^ plugs. There was a decrease in the level of *Tek* expression in *ENG*^*+*^ retinas compared to the level in WT retinas, although we found no differences in *Cdh5* expression (Supplementary Fig. 4a-b).

**Fig. 3.**
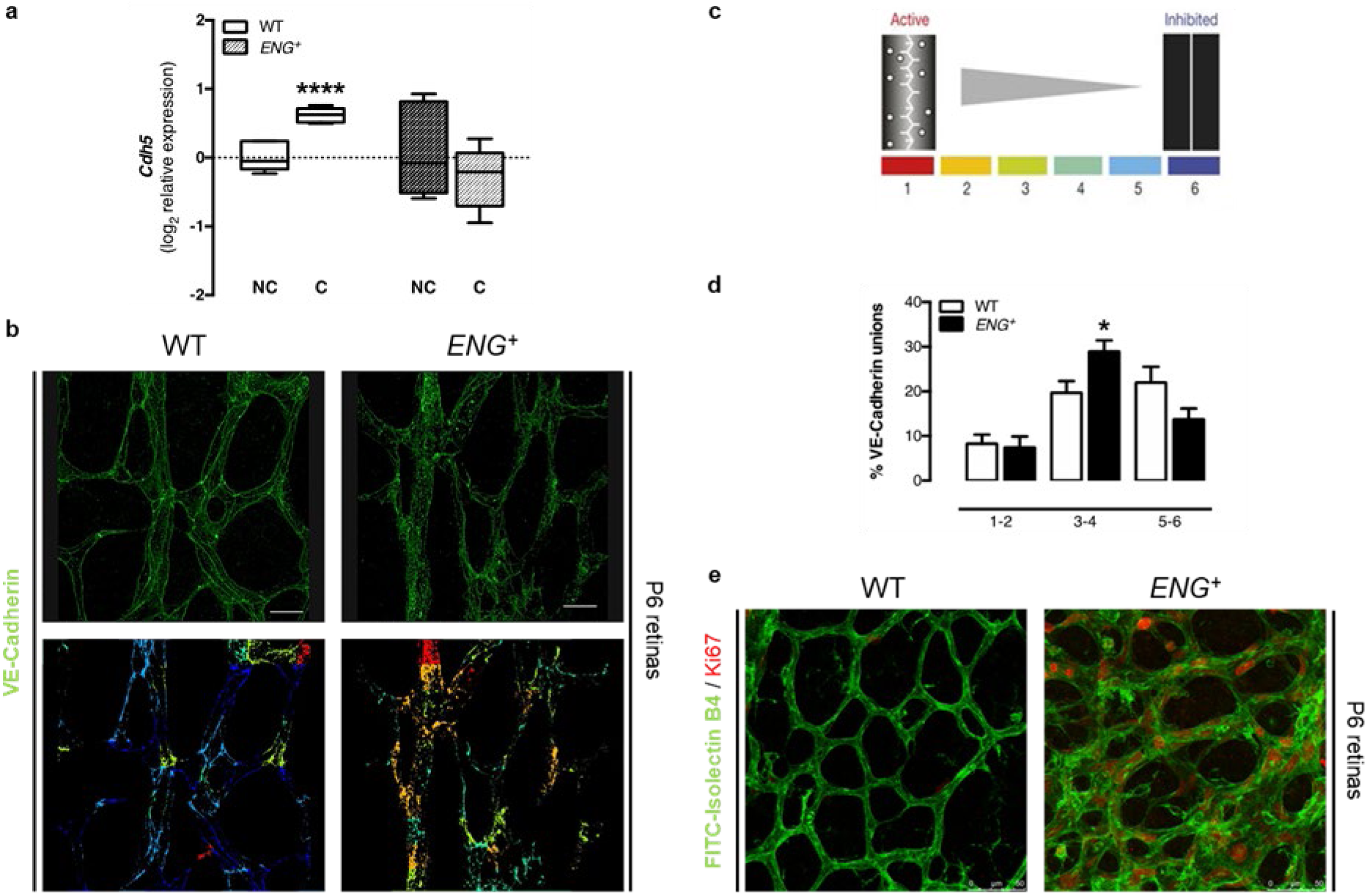
Continuous endoglin overexpression alters endothelium stability *in vivo*. (a) qPCR analysis of *Cdh5* expression in confluent (C) and nonconfluent (NC) MLECs [n(WT)=6, n(*ENG*^*+*^)=6; p=0.0237]. (b) Upper panel: VE-cadherin staining in the retinal vasculature of P6 pups. Lower panel: Pattern of VE-cadherin junctions in the retinal vasculature of P6 pups, using the “patch algorithm” in MATLAB™. (c) Schematic illustration of patch classification numbers and colors used to quantify the pattern of VE-cadherin junctions. (d) Quantification of each type of junction [n(WT)=13, n(*ENG*^*+*^)=12; p (1-2)=0.7905, p(3-4)=0.0183, p(5-6)=0.0732]. (e) Ki67 (red) and FITC-lectin (green) staining in the central area of the retinal vasculature of P6 pups.

However, the role of VE-cadherin in endothelium stabilization depends not only on its expression but also on its distribution along the cell membrane. After the first contacts between ECs, a punctuated pattern of VE-cadherin can be observed. These junctions are called discontinuous, irregular or active and characterize the response to stimuli that reduce the integrity of the endothelial barrier. Subsequently, these complexes create the continuous, regular or inactive adhesions that are present in quiescent vessels (31). For this reason, we analyzed VE-cadherin distribution by immunofluorescence in the central area of the retinal vasculature because under physiological conditions, that should be a stabilized area (Fig. 3b). The image quantification using the “patching algorithm” for MATLAB software (32) demonstrated that compared with WT retinas, *ENG*^*+*^ retinas have a greater number of irregular junctions (Fig. 3b-d). Moreover, immunofluorescence of the proliferation marker Ki67 showed that compared with WT retinas, the central area of the *ENG*^*+*^ retinal vascular plexus contains more proliferating cells (Fig. 3e). These results confirm that continuous endoglin overexpression prevents endothelium stabilization.

### Continuous overexpression of endoglin inhibits pericyte recruitment to the newly formed capillaries

Endoglin has been reported to play a major role in vessel maturation (33,34). Additionally, it has been shown that the RGD sequence of the extracellular domain of endoglin mediates pericyte binding to the ECs (35). However, the greater activation of the endothelium caused by endoglin overexpression could impair mural cell attachment. Thus, we first studied *in vitro* mural cell recruitment in a coculture of EA.hy926 ECs and human brain vascular pericytes (HBVPs) in Matrigel^®^. As discussed above, ECs create capillary-like structures when cultured under these conditions. When cocultured with HBVPs, we observed that compared with Mock cells, *ENG*^*+*^ EA.hy926 ECs recruit fewer mural cells to these structures. Moreover, we studied the adhesion of HBVPs to a monolayer of *ENG*^*+*^ or Mock EA.hy926 ECs, and we found reduced adhesion to ECs that overexpress endoglin (Fig. 4a-c).

**Fig. 4.**
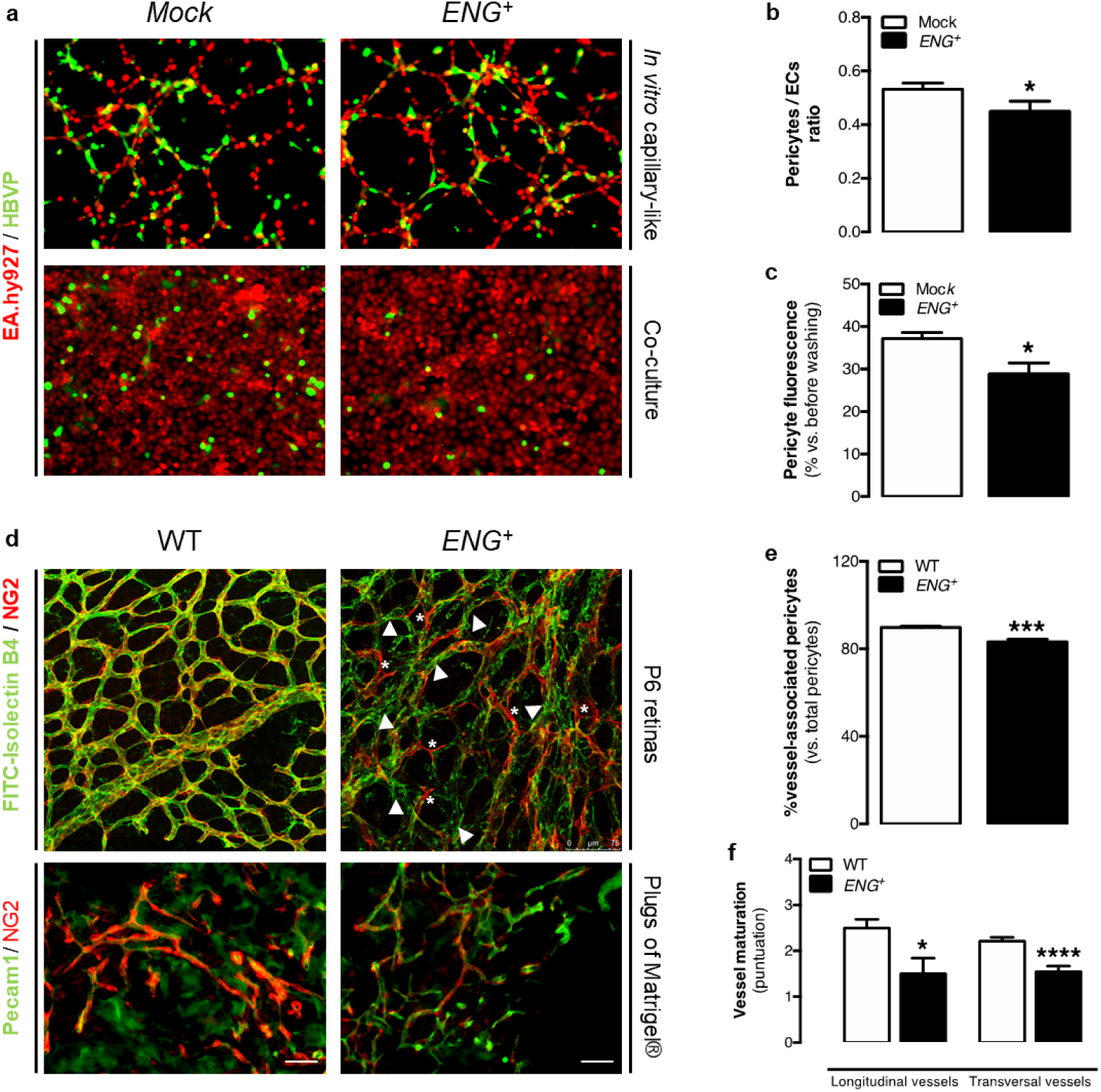
Continuous endoglin overexpression impairs pericyte recruitment *in vivo* and *in vitro*. (a) Upper panel: Pseudocapillary-like structures formed by EA.hy926 cells cocultured with HBVPs in Matrigel^®^. Lower panel: HBVP attachment to EA.hy926 monolayers in culture. (b) Quantification of the ratio between HBVPs and ECs in pseudocapillary-like structures [n(Mock)=3, n(*ENG*^*+*^)=3; p=0.0433]. (c) Quantification of the HBVP fluorescent signal over EA.hy926 monolayers [n(Mock)=3, n(*ENG*^*+*^)=3; p=0.0128]. (d) Upper panel: NG2 (red) and FITC-lectin (green) staining in the retinal vasculature of P6 pups, showing merged signals (yellow) in WT retinas and noncovered endothelium (arrowheads) and mural cells not bound to vessels (asterisk) in *ENG*^*+*^ retinas. Lower panel: NG2 (red) and CD31 (green) staining in plugs of Matrigel^®^ (e) Quantification of the ratio of pericytes that are bound to the endothelium with respect to the total number of pericytes in the retinal vasculature of P6 pups [n(WT)=4, n(*ENG*^*+*^)=6; p<0.0001]. (f) Quantification of the maturation of longitudinal and transversal vessels in plugs of Matrigel^®^, where 1 represents less mature and 3 represents more mature [n(WT)=4, n(*ENG*^*+*^)=3; p (longitudinal)=0.0446, p (transversal)<0.0001].

We also aimed to confirm whether endoglin overexpression decreases vessel maturation *in vivo*. We found no differences in the expression of *Pdgfrb* in retinas from WT and *ENG*^*+*^ mice (Supplementary Fig. 4c), suggesting an equivalent number of mural cells in the retinas. To analyze the distribution of these mural cells, we performed colabeling of the retinal endothelium with FITC-isolectin B4 and NG2, which is a pericyte marker. There was almost complete colocalization in WT retinas, but we observed endothelium that was not entirely covered by mural cells and pericytes that did not appear to be bound to the endothelium in *ENG*^*+*^ retinas, suggesting impaired mural cell attachment (Fig. 4d-e). Similarly, we analyzed the expression of mural cell markers in plugs of Matrigel^®^, and we observed the same results as in retinas. It seems that there are equal numbers of mural cells in plugs (Supplementary Fig. 4d). We then analyzed the vessel maturation state by quantifying the immunofluorescence of the endothelial marker Pecam1 and the mural marker NG2. This assay confirmed that vessels from *ENG*^*+*^ plugs have less mural coverage than vessels from WT plugs (Fig. 4d-f).

### Continuous endoglin overexpression produces leaky vessels that facilitate tumor cell intravasation and metastasis generation

The lower stability of the endothelium and the impaired vessel maturation can lead to an enhanced permeability of the vessels. Therefore, the evidence gathered so far may explain the increased infiltration of erythrocytes in the stroma of tumors developed in *ENG*^*+*^ mice (Fig. 1c). Interestingly, all the vessel structure alterations that we observed after tumor angiogenesis could have more serious consequences, such as the intravasation of tumor cells. For this reason, we analyzed the presence of metastases in WT and *ENG*^*+*^ mice after tumor induction with LLC cells infected with green fluorescence protein (GFP). Mouse lungs were then analyzed to quantify the number of GFP-positive foci as evidence of the presence of tumor cells after metastasis. In agreement with the previous results, compared with WT lungs, *ENG*^*+*^ lungs had more tumor loci (Fig. 5a-b). Moreover, compared with WT mice, *ENG*^*+*^ mice had a higher number of circulating GFP-LLC cells, as evidenced by the fact that blood from these mice generated more GFP^+^ cell colonies *in vitro* (Fig. 5c). These results suggest that continuous endoglin overexpression promotes tumor cell intravasation and the development of metastases in mice.

**Fig. 5.**
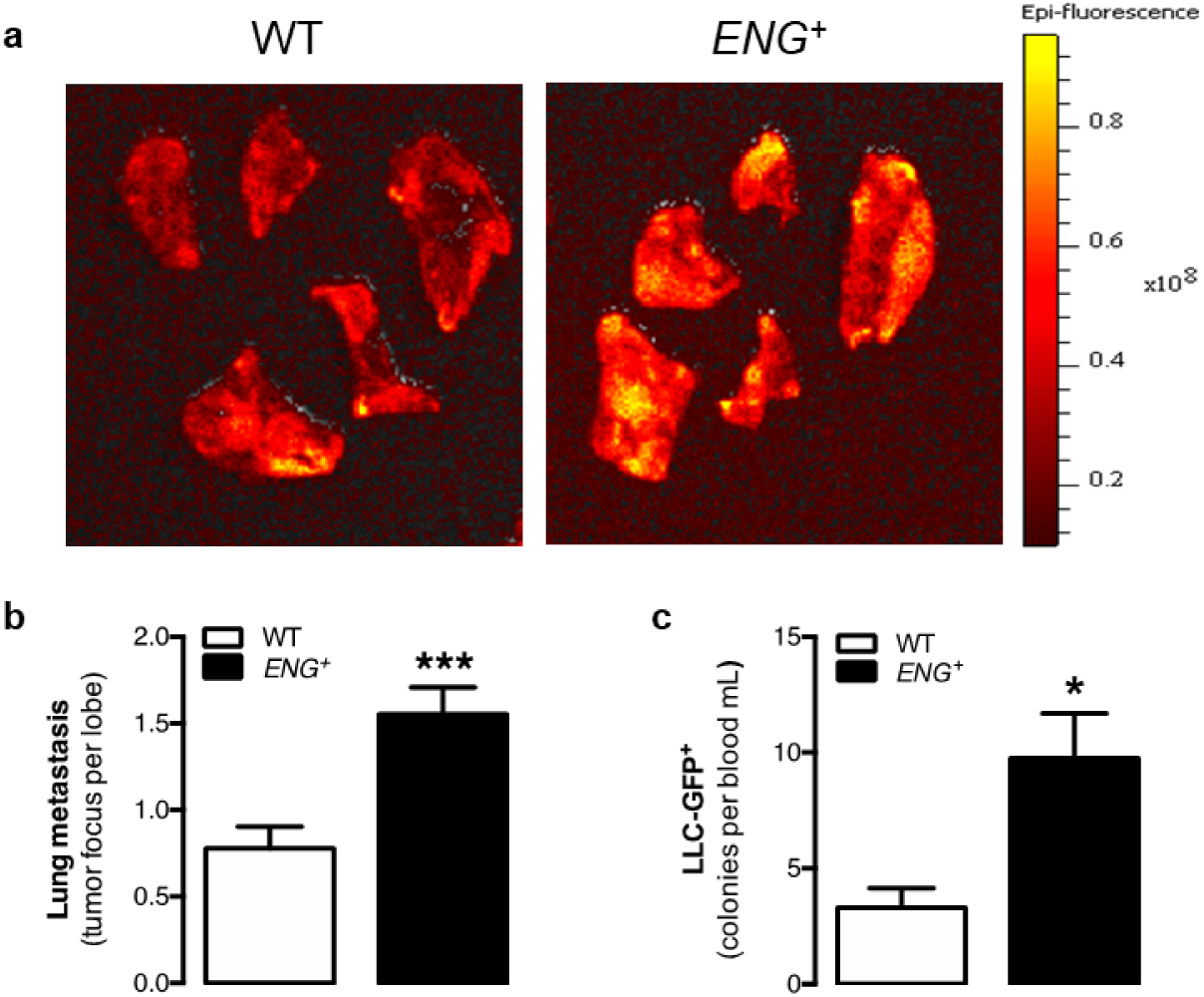
Permanent endoglin overexpression facilitates tumor cell metastasis. (a) Epi-fluorescence images of mouse lung lobe metastases of LLC-GFP^+^ tumor cells. (b) Quantification of tumor metastatic foci per mouse lung lobe [n(WT)=8, n(*ENG*^*+*^)=9; p=0.0004]. (c) Quantification of circulating LLC-GFP^+^ tumor cells in mice [n(WT)=8, n(*ENG*^*+*^)=9; p=0.0107].

## DISCUSSION

In this study, we demonstrated that high levels of endoglin do not improve angiogenesis; instead, constitutive endoglin overexpression produces alterations in the structure of the vasculature. These alterations do not seem to be due to defects during the first phases of angiogenesis; instead, endoglin overexpression prevents vessel stabilization and maturation, as shown *in vitro* and *in vivo*. In tumors, this instability leads to the extravasation of erythrocytes to the tumor stroma and the intravasation of malignant cells, without affecting the tumor size. All this may explain the mechanism, which has never been demonstrated before, by which high levels of endoglin in different solid tumors are associated with a worse prognosis (17,18) and the mechanism by which anti-endoglin therapy reduces the generation of metastases (22,23).

Different authors have proposed that because endoglin deficiency impairs angiogenesis (8,11,13), its pathological overexpression may enhance angiogenesis. However, our results indicate that endoglin overexpression produces a pathological phenotype that affects mainly the stabilization and maturation phases of the angiogenic process. Thus, considering the results of previous studies and our results, we propose a model that could explain the consequences of endoglin levels being above or below physiological levels (Fig. 6a) and the importance of fine-tuned endoglin regulation during angiogenesis. According to this model, it is necessary for endoglin expression to increase during sprouting (8,9) up to a threshold level, which may not be reached in *Eng*^−*/*−^ and *Eng*^*+/*−^ mice, in which the initial phases of angiogenesis do not occur correctly. This is supported by the results of numerous studies that have demonstrated that endoglin expression is increased in the angiogenic edge (8,9) and that its deficiency can alter the proliferation of ECs (8,36,37), migration of ECs (8,11,38) and three-dimensional organization of ECs in Matrigel^®^ (8,11). These alterations may lead to impaired sprouting from *Eng*^*+/*−^ aortic rings (8) and finally to defects in vessel formation in *Eng*^*+/*−^ animals (8,11,13). On the other hand, our results suggest that physiological endoglin expression is enough to develop the initial phases of angiogenesis, as endoglin overexpression does not enhance these phases. In contrast, constitutive endoglin overexpression alters endothelium stabilization and mural cell recruitment, as demonstrated in this study both *in vitro* and *in vivo*. Thus, during the final phases, it seems necessary for endoglin levels to decrease to basal levels to allow the maturation of the vessels, as sustained endoglin overexpression maintains the activation of ECs. Some authors have shown that endoglin-deficient vessels also present altered stabilization and maturation (35,39), giving rise to more permeable vessels (35,40,41). However, this phenotype seems to be explained by the lack of direct interaction between endothelial endoglin and the integrins of pericytes (35).

**Fig. 6.**
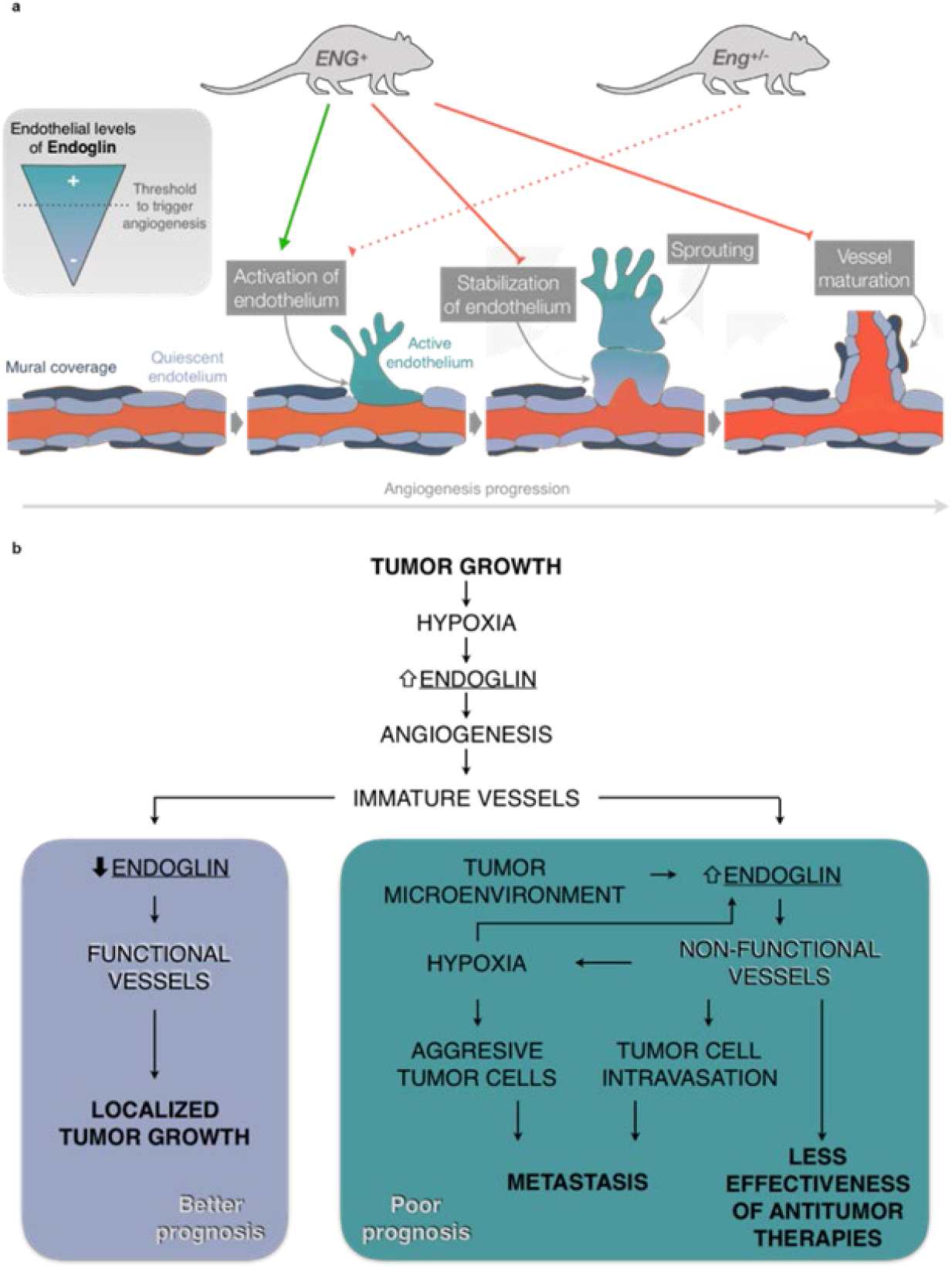
Models of endothelial endoglin regulation during physiological and tumor angiogenesis. (a) Endoglin levels in the endothelium need to be carefully regulated for the correct activation (upregulation) and subsequent stabilization (downregulation to basal levels) of the newly formed vessel. Consistent endoglin overexpression (*ENG*^*+*^) promotes endothelium activation but impedes vessel stabilization and maturation. A lack of endoglin (*Eng*^*+/*−^) may prevent proper activation of the endothelium and triggering of angiogenesis. (b) Tumoral continuous endoglin overexpression could impede vessel normalization which reduces the effectiveness of anti-tumor therapies, enhances the aggressiveness of tumor cells by further increasing hypoxia and promotes the appearance of metastases. On the contrary, downregulation of endoglin after tumor angiogenesis promotes vessel stabilization and thus tumor growth control, which could explain the positive effect of anti-endoglin therapies in cancer control.

Regarding cancer development, other authors have shown that endoglin deficiency or different anti-endoglin treatments reduce the vascularity of tumors and, therefore, decrease tumor growth (21,42–45). However, according to our model and our results, the poor prognosis associated with high levels of endoglin is not related to a greater number of vessels or a larger tumor size. Instead, we propose a second model (Fig. 6b) in which the lack of downregulation of endoglin levels in the tumor endothelium after angiogenesis leads to the lack of stabilization and increased impairment of the function of tumor vessels. This could result in an inadequate nourishment of the tumor, limiting its growth but also increasing hypoxia, which causes tumor cells to acquire a more aggressive phenotype (46,47). In addition, the intravasation of malignant cells could be facilitated by increased vessel instability and impaired mural coverage (48). Accordingly, our results show increased tumor cell intravasation in *ENG*^*+*^ mice compared with WT mice. Considering that metastases are responsible for 90% of cancer deaths, we propose the following new hypothesis: the worse prognosis of patients with tumors with high levels of endoglin is not due to increased tumor size; instead, the overexpression of endoglin maintains the activation of the endothelium and increases the risk of metastases. This hypothesis is in accordance with previous observations of increased metastases in tumors with high levels of endoglin staining (49).

As stated previously, some studies have shown that the administration of anti-endoglin antibodies, such as TRC105, decreases the number of metastases (22,23), although an explanation of the mechanisms by which endoglin contributes to metastasis generation has been lacking. Considering our model (Fig 6b), we hypothesize that anti-endoglin therapies are not only antiangiogenic, as described above (21,50), but also reduce the generation of metastases by promoting the stabilization of the endothelium and the normalization of vessels. One limitation of our study is that endoglin overexpression is ubiquitous, which means that endoglin levels are above the physiological level in all cell types other than ECs, including those in the tumor microenvironment, which may also play an important role in the malignancy of tumors. For example, beyond its antiangiogenic effect on the endothelium, anti-endoglin treatment inhibits cancer-associated fibroblast (CAF) invasion and metastasis (51). However, our *in vitro* results show that endoglin overexpression results in a permanent activation of the endothelium that prevents the maturation of the vessels. Therefore, although it may not be the only cell type involved, we can conclude that the persistence of active endothelium is one of the main causes of the intravasation of tumor cells and the generation of metastases.

Moreover, several studies have shown that the low functionality of tumor vessels prevents conventional anti-tumor therapies from reaching the entire tumor, reducing their effectiveness (46). This is also likely to occur in tumors with high levels of endoglin and may also be related to a worse prognosis (Fig. 6b).

In summary, we demonstrate that, although an increase in endoglin expression is necessary during the first phases of angiogenesis, endoglin expression must return to basal levels to allow mural cell recruitment. This is a key event in tumors, in which the proangiogenic stimuli are very potent and vessel instability facilitates the intravasation of malignant cells and hampers anti-tumor therapy efficacy. Therefore, the normalization of the vasculature resulting from the administration of anti-endoglin therapies may reduce tumor cell intravasation and increase the effectiveness of chemotherapy, as has already been observed with anti-VEGF and ALK1-Fc treatments (46,52,53).

## ACKNOWLEDGMENTS

We thank Carmelo Bernabéu for his excellent suggestions regarding the design of this work and for revising the manuscript; Elena Llano Cuadra for performing the stable infection of EA.hy926 and 3LL cells; Francisco Sánchez-Madrid for generously giving us the anti-endoglin hybridoma; Jesús García Briñón and Javier Herrero Turrión for helping us with the cryostat and confocal microscope, respectively; and Katie Bentley for generously giving us the patching algorithm.

## AUTHOR CONTRIBUTIONS

CO-I and EN-G contributed equally to the design of the work and performed most of the experiments, both *in vitro* and *in vivo*. They also contributed to the writing of the manuscript. CE-T contributed to the metastasis studies and revised the manuscript. LS-S contributed to the immunofluorescence studies and revised the manuscript. AR-B, JML-N and MP supervised the study and contributed to the writing of the manuscript.

## REFERENCES

1. Muz B, de la Puente P, Azab F, Azab AK. The role of hypoxia in cancer progression, angiogenesis, metastasis, and resistance to therapy. Hypoxia [Internet]. 2015 Dec;3:83–92. Available from: https://www.dovepress.com/the-role-of-hypoxia-in-cancer-progression-angiogenesis-metastasis-and--peer-reviewed-article-HP

2. Sherwood LM, Parris EE, Folkman J. Tumor Angiogenesis: Therapeutic Implications. N Engl J Med [Internet]. 1971 Nov 18;285(21):1182–6. Available from: http://www.nejm.org/doi/abs/10.1056/NEJM197111182852108

3. Núñez-Gómez E, Pericacho M, Ollauri-Ibáñez C, Bernabéu C, López-Novoa JM. The role of endoglin in post-ischemic revascularization. Angiogenesis [Internet]. 2017 Feb 9;20(1):1–24. Available from: http://link.springer.com/10.1007/s10456-016-9535-4

4. Eilken HM, Adams RH. Dynamics of endothelial cell behavior in sprouting angiogenesis. Curr Opin Cell Biol [Internet]. 2010 Oct;22(5):617–25. Available from: https://linkinghub.elsevier.com/retrieve/pii/S095506741000133X

5. Geudens I, Gerhardt H. Coordinating cell behaviour during blood vessel formation. Development [Internet]. 2011 Nov 1;138(21):4569–83. Available from: http://dev.biologists.org/cgi/doi/10.1242/dev.062323

6. Carmeliet P, Jain RK. Molecular mechanisms and clinical applications of angiogenesis. Nature [Internet]. 2011 May 19;473(7347):298–307. Available from: http://www.nature.com/articles/nature10144

7. Bernabeu C, Lopez-Novoa JM, Quintanilla M. The emerging role of TGF-β superfamily coreceptors in cancer. Biochim Biophys Acta - Mol Basis Dis [Internet]. 2009 Oct;1792(10):954–73. Available from: https://linkinghub.elsevier.com/retrieve/pii/S0925443909001471

8. Park S, DiMaio TA, Liu W, Wang S, Sorenson CM, Sheibani N. Endoglin regulates the activation and quiescence of endothelium by participating in canonical and non-canonical TGF- signaling pathways. J Cell Sci [Internet]. 2013 Mar 15;126(6):1392–405. Available from: http://jcs.biologists.org/cgi/doi/10.1242/jcs.117275

9. Barnett JM, Suarez S, McCollum GW, Penn JS. Endoglin Promotes Angiogenesis in Cell- and Animal-Based Models of Retinal Neovascularization. Investig Opthalmology Vis Sci [Internet]. 2014 Oct 14;55(10):6490. Available from: http://iovs.arvojournals.org/article.aspx?doi=10.1167/iovs.14-14945

10. McAllister KA, Grogg KM, Johnson DW, Gallione CJ, Baldwin MA, Jackson CE, et al. Endoglin, a TGF-β binding protein of endothelial cells, is the gene for hereditary haemorrhagic telangiectasia type 1. Nat Genet [Internet]. 1994 Dec;8(4):345–51. Available from: http://www.nature.com/articles/ng1294-345

11. Jerkic M, Rodriguezbarbero A, Prieto M, Toporsian M, Pericacho M, Rivaselena J, et al. Reduced angiogenic responses in adult endoglin heterozygous mice. Cardiovasc Res [Internet]. 2006 Mar 1;69(4):845–54. Available from: https://academic.oup.com/cardiovascres/article-lookup/doi/10.1016/j.cardiores.2005.11.020

12. Mahmoud M, Allinson KR, Zhai Z, Oakenfull R, Ghandi P, Adams RH, et al. Pathogenesis of Arteriovenous Malformations in the Absence of Endoglin. Circ Res [Internet]. 2010 Apr 30;106(8):1425–33. Available from: https://www.ahajournals.org/doi/10.1161/CIRCRESAHA.109.211037

13. Sugden WW, Meissner R, Aegerter-Wilmsen T, Tsaryk R, Leonard E V., Bussmann J, et al. Endoglin controls blood vessel diameter through endothelial cell shape changes in response to haemodynamic cues. Nat Cell Biol [Internet]. 2017 May 22;19(6):653–65. Available from: http://www.nature.com/doifinder/10.1038/ncb3528

14. Tien YW, Chang KJ, Jeng YM, Lee PH, Wu MS, Lin JT, et al. Tumor angiogenesis and its possible role in intravasation of colorectal epithelial cells. Clin Cancer Res [Internet]. 2001 Jun;7(6):1627–32. Available from: https://www.ncbi.nlm.nih.gov/pubmed/11410499

15. Basilio-de-Oliveira RP, Pannain VLN. Prognostic angiogenic markers (endoglin, VEGF, CD31) and tumor cell proliferation (Ki67) for gastrointestinal stromal tumors. World J Gastroenterol [Internet]. 2015 Jun 14;21(22):6924–30. Available from: http://www.wjgnet.com/1007-9327/full/v21/i22/6924.htm

16. Bauman TM, Huang W, Lee MH, Abel EJ. Neovascularity as a prognostic marker in renal cell carcinoma. Hum Pathol [Internet]. 2016 Nov 1 [cited 2018 Sep 12];57:98–105. Available from: https://www.sciencedirect.com/science/article/pii/S0046817716301460?via%3Dihub

17. Zhang J, Zhang L, Lin Q, Ren W, Xu G. Prognostic value of endoglin-assessed microvessel density in cancer patients: a systematic review and meta-analysis. Oncotarget [Internet]. 2018 Jan 26;9(7):7660–71. Available from: http://www.oncotarget.com/fulltext/23546

18. Yao Y, Kubota T, Takeuchi H, Sato K. Prognostic significance of microvessel density determined by an anti-CD105/endoglin monoclonal antibody in astrocytic tumors: Comparison with an anti-CD31 monoclonal antibody. Neuropathology [Internet]. 2005 Sep;25(3):201–6. Available from: http://doi.wiley.com/10.1111/j.1440-1789.2005.00632.x

19. Miyata Y, Sagara Y, Watanabe S, Asai A, Matsuo T, Ohba K, et al. CD105 is a more appropriate marker for evaluating angiogenesis in urothelial cancer of the upper urinary tract than CD31 or CD34. Virchows Arch [Internet]. 2013 Nov 25;463(5):673–9. Available from: http://link.springer.com/10.1007/s00428-013-1463-8

20. Sakurai T, Okumura H, Matsumoto M, Uchikado Y, Owaki T, Kita Y, et al. Endoglin (CD105) is a useful marker for evaluating microvessel density and predicting prognosis in esophageal squamous cell carcinoma. Anticancer Res [Internet]. 2014 Jul;34(7):3431–8. Available from: http://www.ncbi.nlm.nih.gov/pubmed/24982351

21. Ollauri-Ibáñez C, López-Novoa JM, Pericacho M. Endoglin-based biological therapy in the treatment of angiogenesis-dependent pathologies. Expert Opin Biol Ther [Internet]. 2017 Sep 2;17(9):1053–63. Available from: https://www.tandfonline.com/doi/full/10.1080/14712598.2017.1346607

22. Paauwe M, Heijkants RC, Oudt CH, van Pelt GW, Cui C, Theuer CP, et al. Endoglin targeting inhibits tumor angiogenesis and metastatic spread in breast cancer. Oncogene [Internet]. 2016 Aug 25;35(31):4069–79. Available from: http://www.nature.com/articles/onc2015509

23. Uneda S, Toi H, Tsujie T, Tsujie M, Harada N, Tsai H, et al. Anti-endoglin monoclonal antibodies are effective for suppressing metastasis and the primary tumors by targeting tumor vasculature. Int J Cancer [Internet]. 2009 Sep 15;125(6):1446–53. Available from: http://doi.wiley.com/10.1002/ijc.24482

24. Oujo B, Muñoz-Félix JM, Arévalo M, Núñez-Gómez E, Pérez-Roque L, Pericacho M, et al. L-Endoglin Overexpression Increases Renal Fibrosis after Unilateral Ureteral Obstruction. Warburton D, editor. PLoS One [Internet]. 2014 Oct 14;9(10):e110365. Available from: https://dx.plos.org/10.1371/journal.pone.0110365

25. Fruttiger M. Development of the retinal vasculature. Angiogenesis [Internet]. 2007 Mar 19;10(2):77–88. Available from: http://link.springer.com/10.1007/s10456-007-9065-1

26. Blanco R, Gerhardt H. VEGF and Notch in Tip and Stalk Cell Selection. Cold Spring Harb Perspect Med [Internet]. 2013 Jan 1;3(1):a006569–a006569. Available from: http://perspectivesinmedicine.cshlp.org/lookup/doi/10.1101/cshperspect.a006569

27. Jakobsson L, Franco CA, Bentley K, Collins RT, Ponsioen B, Aspalter IM, et al. Endothelial cells dynamically compete for the tip cell position during angiogenic sprouting. Nat Cell Biol [Internet]. 2010 Oct 26;12(10):943–53. Available from: http://www.nature.com/articles/ncb2103

28. Gavard J. Endothelial permeability and VE-cadherin. Cell Adh Migr [Internet]. 2014 Mar 25;8(2):158–64. Available from: http://www.tandfonline.com/doi/abs/10.4161/cam.29026

29. Eisa-Beygi S, Macdonald RL, Wen X-Y. Regulatory Pathways Affecting Vascular Stabilization via VE-Cadherin Dynamics: Insights from Zebrafish (Danio Rerio). J Cereb Blood Flow Metab [Internet]. 2014 Sep 16;34(9):1430–3. Available from: http://journals.sagepub.com/doi/10.1038/jcbfm.2014.128

30. Felcht M, Luck R, Schering A, Seidel P, Srivastava K, Hu J, et al. Angiopoietin-2 differentially regulates angiogenesis through TIE2 and integrin signaling. J Clin Invest [Internet]. 2012 Jun 1;122(6):1991–2005. Available from: http://www.jci.org/articles/view/58832

31. Millán J, Cain RJ, Reglero-Real N, Bigarella C, Marcos-Ramiro B, Fernández-Martín L, et al. Adherens junctions connect stress fibres between adjacent endothelial cells. BMC Biol [Internet]. 2010 Dec 2;8(1):11. Available from: https://bmcbiol.biomedcentral.com/articles/10.1186/1741-7007-8-11

32. Bentley K, Franco CA, Philippides A, Blanco R, Dierkes M, Gebala V, et al. The role of differential VE-cadherin dynamics in cell rearrangement during angiogenesis. Nat Cell Biol [Internet]. 2014 Apr 23;16(4):309–21. Available from: http://www.nature.com/articles/ncb2926

33. Bourdeau A, Dumont DJ, Letarte M. A murine model of hereditary hemorrhagic telangiectasia. J Clin Invest [Internet]. 1999 Nov 15;104(10):1343–51. Available from: http://www.jci.org/articles/view/8088

34. Li DY. Defective Angiogenesis in Mice Lacking Endoglin. Science (80-) [Internet]. 1999 May 28;284(5419):1534–7. Available from: http://www.sciencemag.org/cgi/doi/10.1126/science.284.5419.1534

35. Rossi E, Smadja DM, Boscolo E, Langa C, Arevalo MA, Pericacho M, et al. Endoglin regulates mural cell adhesion in the circulatory system. Cell Mol Life Sci [Internet]. 2016 Apr 8;73(8):1715–39. Available from: http://link.springer.com/10.1007/s00018-015-2099-4

36. Pece-Barbara N, Vera S, Kathirkamathamby K, Liebner S, Di Guglielmo GM, Dejana E, et al. Endoglin Null Endothelial Cells Proliferate Faster and Are More Responsive to Transforming Growth Factor β1 with Higher Affinity Receptors and an Activated Alk1 Pathway. J Biol Chem [Internet]. 2005 Jul 29;280(30):27800–8. Available from: http://www.jbc.org/lookup/doi/10.1074/jbc.M503471200

37. Pan CC, Kumar S, Shah N, Hoyt DG, Hawinkels LJAC, Mythreye K, et al. Src-mediated Post-translational Regulation of Endoglin Stability and Function Is Critical for Angiogenesis. J Biol Chem [Internet]. 2014 Sep 12;289(37):25486–96. Available from: http://www.jbc.org/lookup/doi/10.1074/jbc.M114.578609

38. Lee NY, Blobe GC. The Interaction of Endoglin with β-Arrestin2 Regulates Transforming Growth Factor-β-mediated ERK Activation and Migration in Endothelial Cells. J Biol Chem [Internet]. 2007 Jul 20;282(29):21507–17. Available from: http://www.jbc.org/lookup/doi/10.1074/jbc.M700176200

39. Lebrin F, Srun S, Raymond K, Martin S, van den Brink S, Freitas C, et al. Thalidomide stimulates vessel maturation and reduces epistaxis in individuals with hereditary hemorrhagic telangiectasia. Nat Med [Internet]. 2010 Apr 4;16(4):420–8. Available from: http://www.nature.com/articles/nm.2131

40. Anderberg C, Cunha SI, Zhai Z, Cortez E, Pardali E, Johnson JR, et al. Deficiency for endoglin in tumor vasculature weakens the endothelial barrier to metastatic dissemination. J Exp Med [Internet]. 2013 Mar 11;210(3):563–79. Available from: http://www.ncbi.nlm.nih.gov/pubmed/23401487

41. Jerkic M, Letarte M. Increased endothelial cell permeability in endoglin-deficient cells. FASEB J [Internet]. 2015 Sep;29(9):3678–88. Available from: http://www.fasebj.org/doi/10.1096/fj.14-269258

42. D&uuml;wel A, Eleno N, Jerkic M, Arevalo M, Bola&ntilde;os JP, Bernabeu C, et al. Reduced Tumor Growth and Angiogenesis in Endoglin-Haploinsufficient Mice. Tumor Biol [Internet]. 2007;28(1):1–8. Available from: http://www.karger.com/doi/10.1159/000097040

43. Dolinsek T, Markelc B, Sersa G, Coer A, Stimac M, Lavrencak J, et al. Multiple Delivery of siRNA against Endoglin into Murine Mammary Adenocarcinoma Prevents Angiogenesis and Delays Tumor Growth. Pizzo S V., editor. PLoS One [Internet]. 2013 Mar 5;8(3):e58723. Available from: https://dx.plos.org/10.1371/journal.pone.0058723

44. Dolinsek T, Markelc B, Bosnjak M, Blagus T, Prosen L, Kranjc S, et al. Endoglin Silencing has Significant Antitumor Effect on Murine Mammary Adenocarcinoma Mediated by Vascular Targeted Effect. Curr Gene Ther [Internet]. 2015 Mar 28;15(3):228–44. Available from: http://www.eurekaselect.com/openurl/content.php?genre=article&issn=1566-5232&volume=15&issue=3&spage=228

45. Stimac M, Kamensek U, Cemazar M, Kranjc S, Coer A, Sersa G. Tumor radiosensitization by gene therapy against endoglin. Cancer Gene Ther [Internet]. 2016 Jul 20;23(7):214–20. Available from: http://www.nature.com/articles/cgt201620

46. Jain RK, Martin JD, Stylianopoulos T. The Role of Mechanical Forces in Tumor Growth and Therapy. Annu Rev Biomed Eng [Internet]. 2014 Jul 11;16(1):321–46. Available from: http://www.ncbi.nlm.nih.gov/pubmed/25014786

47. Reymond N, D’Água BB, Ridley AJ. Crossing the endothelial barrier during metastasis. Nat Rev Cancer [Internet]. 2013 Dec 1;13(12):858–70. Available 46. from: http://www.nature.com/articles/nrc3628

48. García-Román J, Zentella-Dehesa A. Vascular permeability changes involved in tumor metastasis. Cancer Lett [Internet]. 2013 Jul;335(2):259–69. Available from: https://linkinghub.elsevier.com/retrieve/pii/S0304383513002280

49. Chien C-Y, Su C-Y, Hwang C-F, Chuang H-C, Chen C-M, Huang C-C. High expressions of CD105 and VEGF in early oral cancer predict potential cervical metastasis. J Surg Oncol [Internet]. 2006 Oct 1;94(5):413–7. Available from: http://doi.wiley.com/10.1002/jso.20546

50. Liu Y, Tian H, Blobe GC, Theuer CP, Hurwitz HI, Nixon AB. Effects of the combination of TRC105 and bevacizumab on endothelial cell biology. Invest New Drugs [Internet]. 2014 Oct 5;32(5):851–9. Available from: http://link.springer.com/10.1007/s10637-014-0129-y

51. Paauwe M, Schoonderwoerd MJ, Helderman RFCPA, Harryvan TJ, Groenewoud A, van Pelt GW, et al. Endoglin Expression on Cancer-Associated Fibroblasts Regulates Invasion and Stimulates Colorectal Cancer Metastasis. Clin Cancer Res [Internet]. 2018 Dec 15;24(24):6331–44. Available from: http://clincancerres.aacrjournals.org/lookup/doi/10.1158/1078-0432.CCR-18-0329

52. Hawinkels LJAC, de Vinuesa AG, Paauwe M, Kruithof-de Julio M, Wiercinska E, Pardali E, et al. Activin Receptor-like Kinase 1 Ligand Trap Reduces Microvascular Density and Improves Chemotherapy Efficiency to Various Solid Tumors. Clin Cancer Res [Internet]. 2016 Jan 1;22(1):96–106. Available from: http://clincancerres.aacrjournals.org/cgi/doi/10.1158/1078-0432.CCR-15-0743

53. Arjaans M, Oude Munnink TH, Oosting SF, Terwisscha van Scheltinga AGT, Gietema JA, Garbacik ET, et al. Bevacizumab-Induced Normalization of Blood Vessels in Tumors Hampers Antibody Uptake. Cancer Res [Internet]. 2013 Jun 1;73(11):3347–55. Available from: http://cancerres.aacrjournals.org/cgi/doi/10.1158/0008-5472.CAN-12-3518

