## Supplementary figures and tables for "Continuous Endoglin (CD105) Overexpression Disrupts Angiogenesis and Facilitates Tumor Cell Metastasis"

### SUPPLEMENTARY DATA

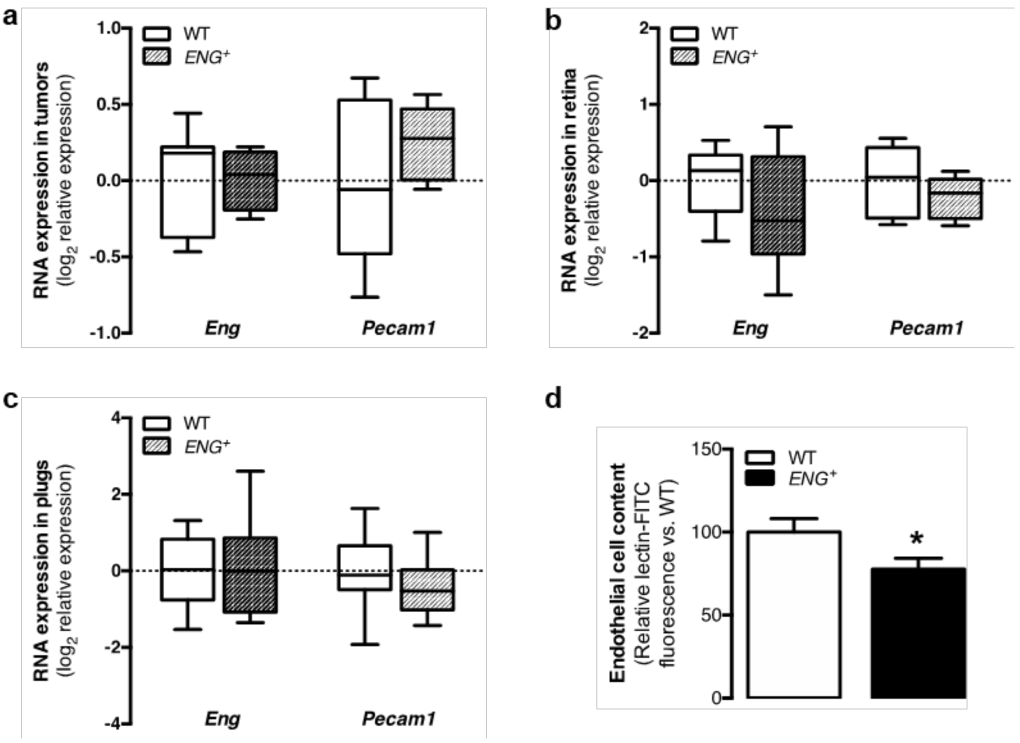


**Supplementary Fig. 1. Continuous endoglin overexpression in tumor and physiological angiogenesis.** (a) qPCR analysis of *Eng* and *Pecam1* expression in LLC tumors [n(WT)=6, n(*ENG*$^+$)=6; p(*Eng*)=0.9657, p(*Pecam1*)=0.2363]. (b) qPCR analysis of *Eng* and *Pecam1* expression in the retinas of P6 mouse pups [n(WT)=6, n(*ENG*$^+$)=6; p(*Eng*)=0.3361, p(*Pecam1*)=0.3785]. (c) qPCR analysis of *Eng* and *Pecam1* expression in plugs of Matrigel® [n(WT)=11, n(*ENG*$^+$)=9; p(*Eng*)=0.8832, p(*Pecam1*)=0.2980]. (d) Quantification of angioreactor EC content, measured by the FITC-lectin signal 9 days after implantation [n(WT)=12, n(*ENG*$^+$)=11; p=0.0479].

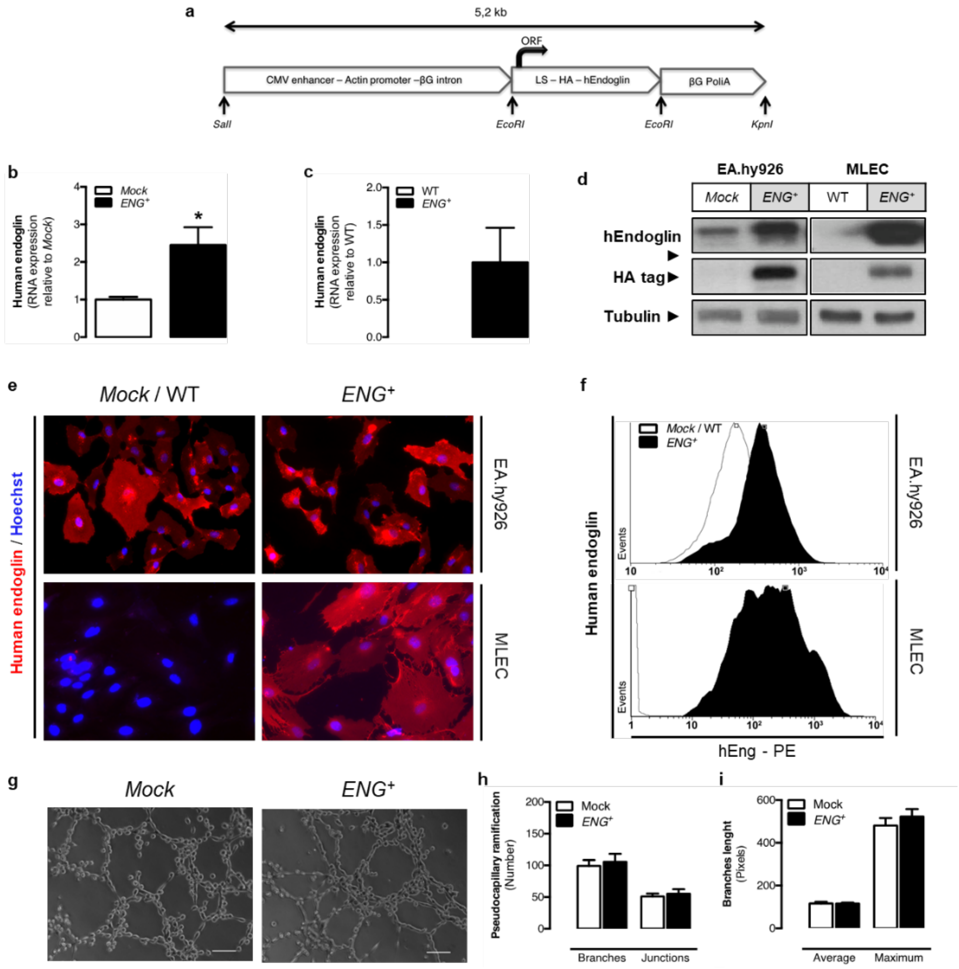


**Supplementary Fig. 2. Characterization of transgenic ECs overexpressing endoglin.** (a) Structure of the vector containing the human endoglin gene used for the stable infection of EA.hy926 cells to generate endoglin overexpression (*ENG*$^{+}$) and *ENG*$^{+}$ mice (MLEC source). The same vector without the endoglin gene was used to generate *Mock* EA.hy926 cells. (b) qPCR detection of human endoglin expression in *Mock* and *ENG*$^{+}$ EA.hy926 cells. (c) qPCR detection of human endoglin expression in WT and *ENG*$^{+}$ MLEC cells. (d) Western blot assessment of human endoglin in the lysates of EA.hy926 and MLEC cells. (e) Immunofluorescence of human endoglin in EA.hy926 and MLEC cells. (f) FACS measurement of surface human endoglin in EA.hy926 and MLEC cells. (g) Pseudocapillary-like structures formed by *Mock* and *ENG*$^{+}$ EA.hy926 cells in Matrigel®. (h) Quantification of the number of branches and junctions of the EA.hy926 pseudocapillary-like structures [n(Mock)=3, n(*ENG*$^{+}$)=3; p(branches)=0.6819, p(junctions)=0.6164]. (i) Quantification of the average and maximum lengths of the branches of the EA.hy926 pseudocapillary-like structures [n(Mock)=3, n(*ENG*$^{+}$)=3; p(average)=0.9536, p(maximum)=0.4029].

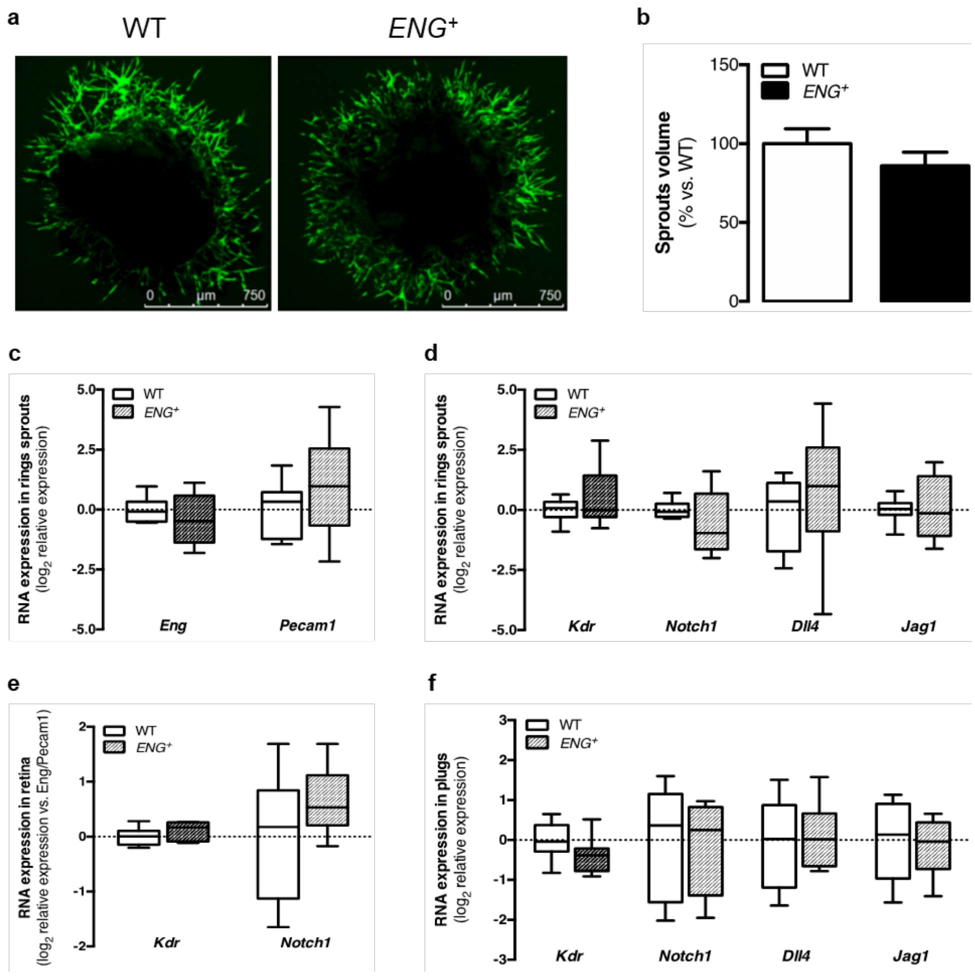


**Supplementary Fig. 3. Sprouting and tip/stalk selection-related gene expression analysis upon endoglin overexpression.** (a) FITC-lectin-labeled sprout growth from aortic rings isolated from WT and *ENG*$^{+}$ mice. (b) Quantification of the volume occupied by sprouts [n(WT)=25, n(*ENG*$^{+}$)=20; p=0.2875]. (c) qPCR analysis of *Eng* and *Pecam1* expression in sprouts from aortic rings [n(WT)=10, n(*ENG*$^{+}$)=10; p(*Eng*)=0.3419, p(*Pecam1*)=0.1960]. (d) qPCR analysis of *Kdr*, *Dll4*, *Notch1* and *Jag1* expression in sprouts from aortic rings [n(WT)=9, n(*ENG*$^{+}$)=9; p(*Kdr*)=0.1787, p(*Dll4*)=0.4884, p(*Notch1*)=0.2344, p(*Jag1*)=0.8735]. (e) qPCR analysis of *Kdr* and *Notch1* expression in the retinas of p6 mouse pups [n(WT)=6, n(*ENG*$^{+}$)=6; p(*Kdr*)=0.2881, p(*Notch1*)=0.2710]. (f) qPCR analysis of *Kdr*, *Dll4*, *Notch1* and *Jag1* expression in plugs of Matrigel® [n(Mock)=9, n(*ENG*$^{+}$)=8; p(*Kdr*)=0.0890, p(*Dll4*)=0.8956, p(*Notch1*)=0.8338, p(*Jag1*)=0.7124].

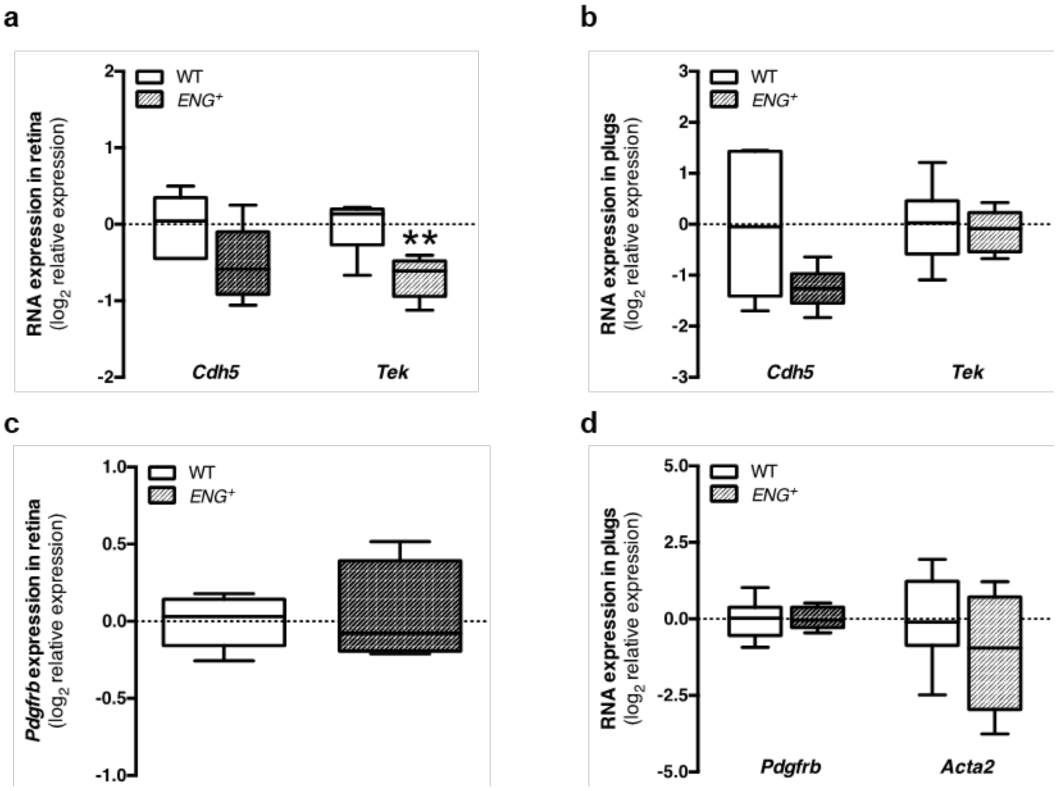


**Supplementary Fig. 4. The effect of permanent endoglin overexpression on endothelium stabilization and mural cell coverage.** (a) qPCR analysis of *Cdh5* and *Tek* expression in the retinas of p6 mouse pups [n(WT)=6, n(*ENG*$^{+}$)=6; p(*Cdh5*)=0.0733, p(*Tek*)=0.0061]. (b) qPCR analysis of *Cdh5* and *Tek* expression in plugs of Matrigel® [n(WT)=9, n(*ENG*$^{+}$)=8; p(*Cdh5*)=0.0684, p(*Tek*)=0.6843]. (c) qPCR analysis of *Pdgfrb* expression in the retinas of p6 mouse pups [n(WT)=6, n(*ENG*$^{+}$)=6; p=0.6977]. (d) qPCR analysis of *Pdgfrb* and *Acta2* expression in plugs of Matrigel® [n(WT)=9, n(*ENG*$^{+}$)=8; p(*Pdgfrb*)=0.9065, p(*Acta2*)=0.2327].

**Table 1. Primers used for cDNA preamplification and qPCR in aortic rings and plugs.**

| Gene | Preamplication primers (Bio-Rad) | qPCR primers (Bio-Rad) |
|---|---|---|
| *Eng* | qMmuCED0046322 | qMmuCID0010792 |
| *Pecam1* | qMmuCID0005317 | qMmuCID0005317 |
| *Dll4* | qMmuCID0016240 | qMmuCID0016240 |
| *Kdr* | qMmuCID005890 | qMmuCID0005890 |
| *Notch1* | qMmuCED0045879 | qMmuCED0045879 |
| *Jag1* | qMmuCID0022326 | qMmuCID0022326 |
| *Tek* | qMmuCID0015486 | qMmuCID0015486 |
| *Cdh5* | qMmuCID0005343 | qMmuCID0005343 |
| *Pdgfrb* | qMmuCED0045914 | qMmuCED0045914 |
| *Acta2* | qMmuCID0006375 | qMmuCID0006375 |

**Table 2. Primer sequences.**

| **Gene** | **qPCR primers** |
|---|---|
| *ENG* | AGGTGCTTCTGGTCCTCAGT<br>CCACTCAAGGATCTGGGTCT |
| *Eng* | GACTTCAGATTGGAATACCTTGG<br>CAGTGCCGTGTCTTTCTGTAAT |
| *Pecam1* | AGGACAACCGTACCTTGGGTGACT<br>CAGTTCTGACACGTACCGGGTCTC |
| *Kdr* | GGCGACTATGTTTGCTCTGC<br>GCCATGCGCTCTAGGATGAT |
| *Notch1* | CAGTGCAACCCCCTGTATGA<br>TCTAGGCCATCCCACTCACA |
| *Tek* | AACAAGAGCGAGTGGACCAT<br>TCCATGGCGCCTTCTACTAC |
| *Cdh5* | ATTGGCCTGTGTTTTCGCAC<br>CACAGTGGGGTCATCTGCAT |
| *Pdgfrb* | AGGACAACCGTACCTTGGGTGACT<br>CAGTTCTGACACGTACCGGGTCTC |
| *Acta2* | AGCCATCTTTCATTGGGATGG<br>CCCCTGACAGGACGTTGTTA |
